# Development of migrating entheses involves replacement of progenitor populations

**DOI:** 10.1101/210799

**Authors:** Neta Felsenthal, Sarah Rubin, Tomer Stern, Sharon Krief, Deepanwita Pal, Brian A. Pryce, Ronen Schweitzer, Elazar Zelzer

## Abstract

Attachment sites of tendons to bones, called entheses, are essential for proper musculoskeletal function. They are formed embryonically by *Sox9*+ progenitors and undergo a developmental process that continues into the postnatal period and involves *Gli1* lineage cells. During bone elongation, some entheses maintain their relative positions by actively migrating along the bone shaft, while others, located at the bone’s extremities, remain stationary. Despite their importance, we lack information on the developmental transition from embryonic to mature enthesis and on the relation between *Sox9*+ progenitors and *Gli1* lineage cells. Here, by performing a series of lineage tracing experiments, we identify the onset of *Gli1* lineage contribution to different entheses during embryogenesis. We show that *Gli1* expression is regulated by SHH signaling during embryonic development, whereas postnatally it is maintained by IHH signaling. Interestingly, we found that unlike in stationary entheses, where *Sox9*+ cells differentiate into the *Gli1* lineage, in migrating entheses the *Sox9* lineage is replaced by *Gli1* lineage and do not contribute to the mature enthesis. Moreover, we show that these *Gli1+* progenitors are pre-specified embryonically to form the different cellular domains of the mature enthesis.

Overall, these findings demonstrate a developmental strategy whereby one progenitor population establishes a simple, embryonic tissue, whereas another population is responsible for its maturation into a complex structure during its migration. Moreover, they suggest that different cell populations may be considered for cell-based therapy of enthesis injuries.

## INTRODUCTION

The proper assembly of the musculoskeletal system is essential for the function, form and stability of the organism. During embryogenesis, an attachment between tendon and bone, known as enthesis, is formed. Thereafter and into the postnatal period, the rudimentary enthesis further develops into a more complex tissue. While some knowledge on enthesis formation and maturation exists, far less is known about the processes that transform the simple embryonic enthesis into the structure of the mature enthesis.

Traditionally, entheses are divided to either fibrous or fibrocartilaginous, according to their composition. Fibrous entheses form at the attachment site of a tendon that is inserted directly into the bone shaft, forming a structure that resembles a root system. This structure, which is composed of a dense connective tissue (Doschak and Zernicke, 2005; Shaw and Benjamin, 2007; Wang et al., 2013), was shown to be regulated by parathyroid hormone-like hormone (PTHLH, also known as PTHrP) (Wang et al., 2013). Fibrocartilage entheses typically form at attachment sites of tendons to the epiphysis or to bone eminences. Relative to fibrous entheses, fibrocartilage entheses are structurally more complex, displaying a cellular gradient that is typically divided into four zones, namely tendon, fibrocartilage, mineralized fibrocartilage and bone. The graded tissue that develops postnatally dissipates the stress that forms at the attachment site, thereby providing the enthesis with the mechanical strength necessary to withstand compression (Benjamin et al., 2006).

Enthesis development is initiated by the specification of a specialized pool of progenitor cells that express both SRY-box 9 (*Sox9*), a key regulator of chondrogenesis, and the tendon marker scleraxis (*Scx*) (Akiyama et al., 2005; Blitz et al., 2013; Schweitzer et al., 2001; Sugimoto et al., 2013). Cell lineage studies showed that *Sox9*-expressing progenitor cells contribute to the formation of the enthesis in neonatal mice (Akiyama et al., 2005; Soeda et al., 2010). Another marker for enthesis cells is GLI-Kruppel family member GLI1 (*Gli1*), a component of the hedgehog (HH) signaling pathway. Lineage studies revealed that *Gli1*-expressing cells act as progenitors that contribute to the formation of some adult entheses (Dyment et al., 2015; Schwartz et al., 2015). However, the relation between progenitors of the embryonic enthesis and *Gli1* lineage cells, which contribute to the mature enthesis, has not been determined.

The relative position of an enthesis along the bone directly affects its mechanical function and, subsequently, the animal’s mobility (Polly, 2007; Salton and Sargis, 2009). Recently, it was shown that all entheses maintain their relative position during bone elongation (Stern et al., 2015). The mechanism that maintains their positions involves regulation of the relative growth rates at the two epiphyseal plates. Additionally, some entheses migrate through continuous reconstruction to maintain their position, a process known as bone modelling or drift (Benjamin and McGonagle, 2009; Dörfl, 1980a; Dörfl, 1980b). These entheses, referred to in the following as migrating entheses, face a unique developmental challenge. Unlike most organs and tissues, they must develop into a complex graded tissue while constantly drifting. This raises the question of whether the descendants of the embryonic enthesis progenitors continue to serve as the building blocks during maturation of migrating entheses.

In this work, we identify the embryonic stage at which *Gli1* lineage is initiated and demonstrate its contribution to the postnatal enthesis. We show that embryonic *Gli1* expression is initially under the regulation of SHH. Later during postnatal development, Gli1 expression is maintained by IHH. Moreover, we show that *Sox9* lineage does not contribute to postnatal migrating entheses. Instead, *Gli1* lineage cells replace the *Sox9* lineage cells and populate the enthesis. Finally, we show that embryonic *Gli1* lineage cells are pre-determined to contribute to the different layers of the fibrocartilaginous enthesis.

## RESULTS

### Some entheses undergo cellular and morphological changes while migrating

The development of the rudimentary embryonic attachment site into the complex structure of a mature enthesis has received little attention (Galatz et al., 2007). As mentioned, in addition to substantial cellular and morphological changes, some entheses also migrate considerably along the bone during bone growth (Benjamin and McGonagle, 2009; Dörfl, 1980a; Dörfl, 1980b; Stern et al., 2015). Therefore, to study the transition that migrating entheses undergo during maturation, we documented morphological and molecular changes as well as drifting activity in entheses from embryonic day (E) 14.5 to postnatal day (P) 14. We focused on two migrating entheses, namely the deltoid enthesis (DT), a fibrocartilaginous enthesis that forms between the deltoid tendon and the deltoid tuberosity, and the teres major enthesis (TM), a fibrous enthesis that forms between the teres major tendon and the humeral shaft. Analysis of enthesis positions during development revealed that both DT (0.778±0.026 mm) and TM (1.624±0.171 mm) entheses drifted considerably along the bone shaft during bone elongation (Fig. 1A). Histological sections through wild-type (WT) mouse humeri showed that both embryonic entheses displayed a simple structure of layered cells (Fig. 1B,C(a),D(a)). However, the overall shape of the enthesis dramatically changed through development, as it protruded outwards from the bone shaft and the different enthesis domains became more noticeable (Fig. 1C(a’-a’’’),D(a’-a’’’). Tissue complexity also increased, as a larger variety of cells, such as fibrocartilage cells and osteoblasts (Benjamin and McGonagle, 2009), were identified along with an increase in extracellular matrix (Fig. 1C(d,d’,h,h’), D(d,d’,h,h’)). The increased complexity was also demonstrated by a change in the expression patterns of structural genes, such as bone sialoprotein (*Bsp*; also known as integrin binding sialoprotein (*Ibsp*), collagen type 2 alpha 1 (*Col2a1*), collagen type 12 alpha 1 (*Col12a1)* and tenascin C (*Tnc*), and regulatory genes such as *Gli1*. Furthermore, expression domains correlating to various structural domains emerged, namely *Col12a1*, *Tnc, Gli1* and *Col1a1* in tendon and fibrocartilage (Fig. 1C(c,f-h),D(c,f-h), and *Col1a1, Gli1* and *Bsp* in mineralized fibrocartilage and bone (Fig. 1C(c,e,h),D(c,e,h)).

**Figure 1.**
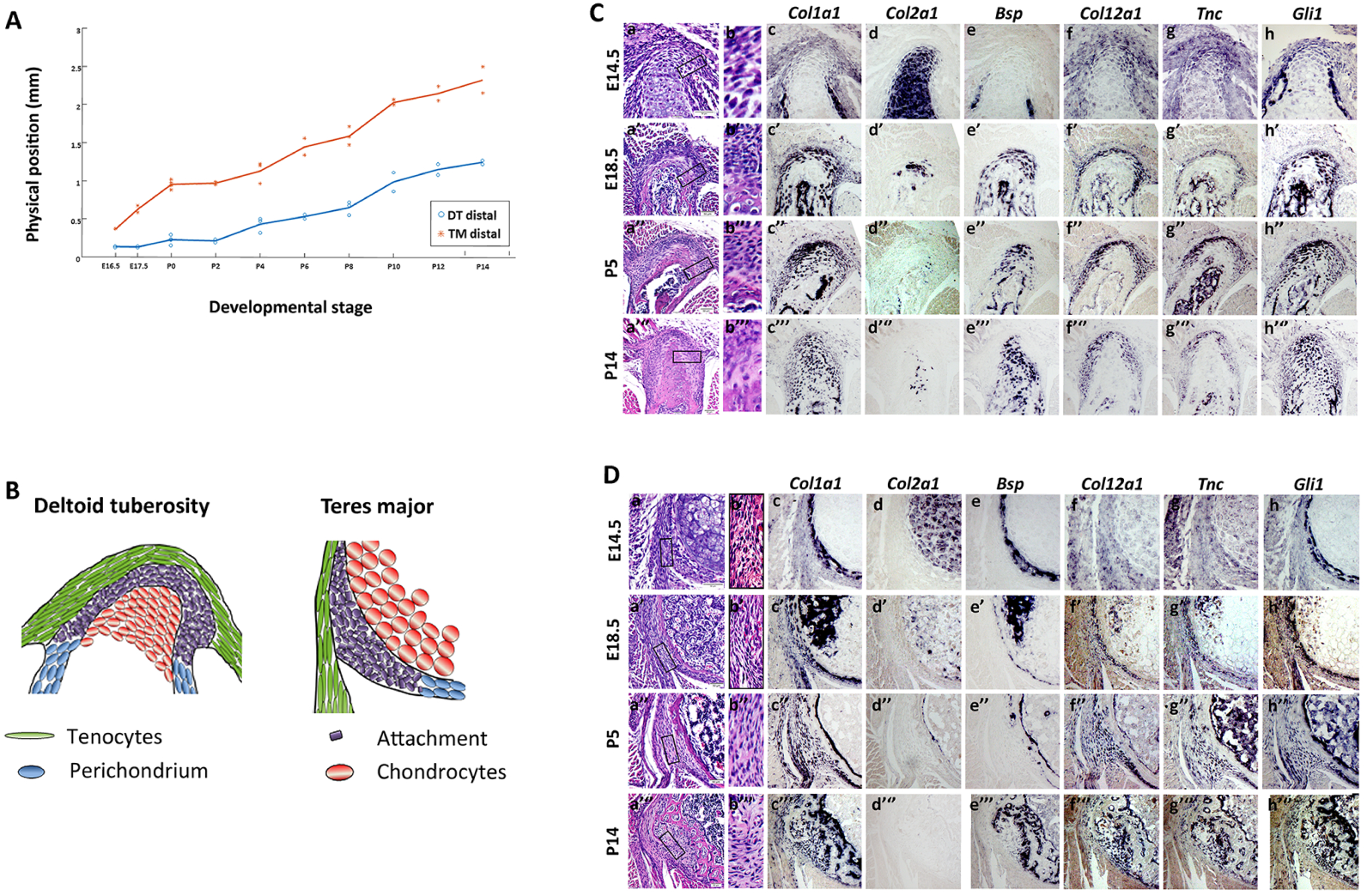
Some entheses undergo cellular and morphological changes while migrating. (**A**) Graph showing the physical position of the deltoid tuberosity (DT) and teres major (TM) entheses along the bone throughout development (E16.5-P14), defined as the distance of the element (mm) from a predefined longitudinal origin roughly at the center of the shaft. Positive and negative values correspond to elements that are proximal and distal to the point of origin, respectively. As indicated by the steep slope of the curves, both DT and TM migrate considerably through development. **(B)** Schematic illustrations of the embryonic DT and TM entheses. **(C and D)** Transverse sections through the humerus showing the DT (A) and TM (B) entheses of E14.5-P14 WT mice. [C(a,b) and D(a,b)]: H…E staining. C(b) and D(b) are magnifications of the boxed areas in C(a) and D(a), respectively. [C(c-h) and D(c-h)]: *In situ* hybridization using digoxigenin-labeled, anti-sense complementary RNA probes for *Col1a1*, *Col2a1*, *Bsp*, *Col12a1*, *Tnc* and *Gli1* at E14.5 (c-h), E18.5 (c’-h’), P5 (c’’-h’’) and P14 (c’’’-h’’’). Scale bars: 50 µm.

### *Gli1*+ cell lineage contributes to the postnatal enthesis

Previously, lineage tracing experiments showed that *Gli1* lineage cells contribute to postnatal enthesis development (Dyment et al., 2015; Schwartz et al., 2015). Yet, the onset of this lineage during embryogenesis and its dynamics in different entheses have been missing. In order to identify the onset of *Gli1* lineage, we performed pulse-chase experiments on *Gli1-CreER*^*T2*^ (Ahn & Joyner, 2004) mice crossed with *R26R-tdTomato* reporter mice (Madisen et al., 2010), which allowed us to mark *Gli1* expressing cells at specific time points and follow their descendants. We analyzed three entheses representing different types, namely DT (migratory-fibrocartilaginous), TM (migratory-fibrous), and Achilles (stationary-fibrocartilaginous). Examination of neonatal (P0) TM and DT entheses following tamoxifen administration at E11.5 and E12.5 revealed only a few *Gli1* lineage cells. However, administration at E13.5 and E15.5 resulted in extensive labeling in both entheses (Fig. 2A-A’’’,B-B’’’). In the stationary Achilles enthesis, extensive labeling was observed at P0 only after tamoxifen administration at E15.5 (Fig. 3C’’’). The postnatal contribution of *Gli1* lineage to migratory entheses was further established by following E13.5 lineage induction to P14 (Fig. 2D,E).

**Figure 2.**
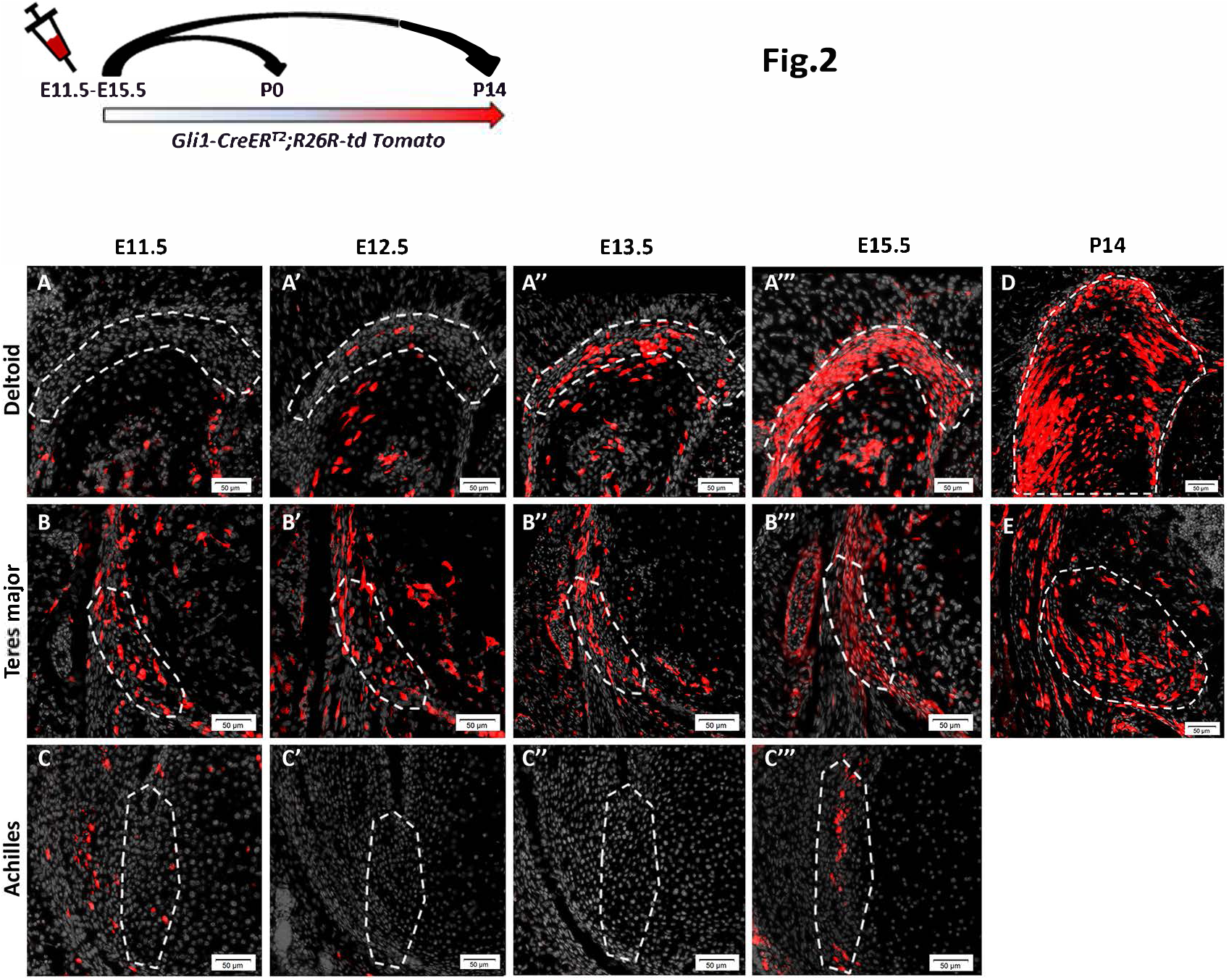
*Gli1+* cell lineage contributes to the postnatal enthesis. Pulse-chase cell lineage experiments using *Gli1-CreER^T2^;R26R-tdTomato* mice demonstrate the contribution of *Gli1* lineage cells to various entheses. *Gli1*-positive cells were pulsed by a single tamoxifen administration at different time points (E11.5-E15.5) and their descendants were followed to P0 and P14. **(A-A’’’)** The DT enthesis was extensively marked after tamoxifen administration at E13.5 and E15.5. **(B-B’’’)** tdTomato-positive cells were identified in the TM enthesis regardless of the time of pulsing; however, a stronger signal was seen following tamoxifen administration at E13.5 or later. **(C-C’’’)** tdTomato-positive cells were identified in the Achilles tendon enthesis following tamoxifen administration at E15.5. **(D-E)** Both DT and TM entheses were extensively marked at P14 following tamoxifen administration at E13.5. The enthesis is demarcated by a dashed line. Scale bars: 50 µm.

**Figure 3.**
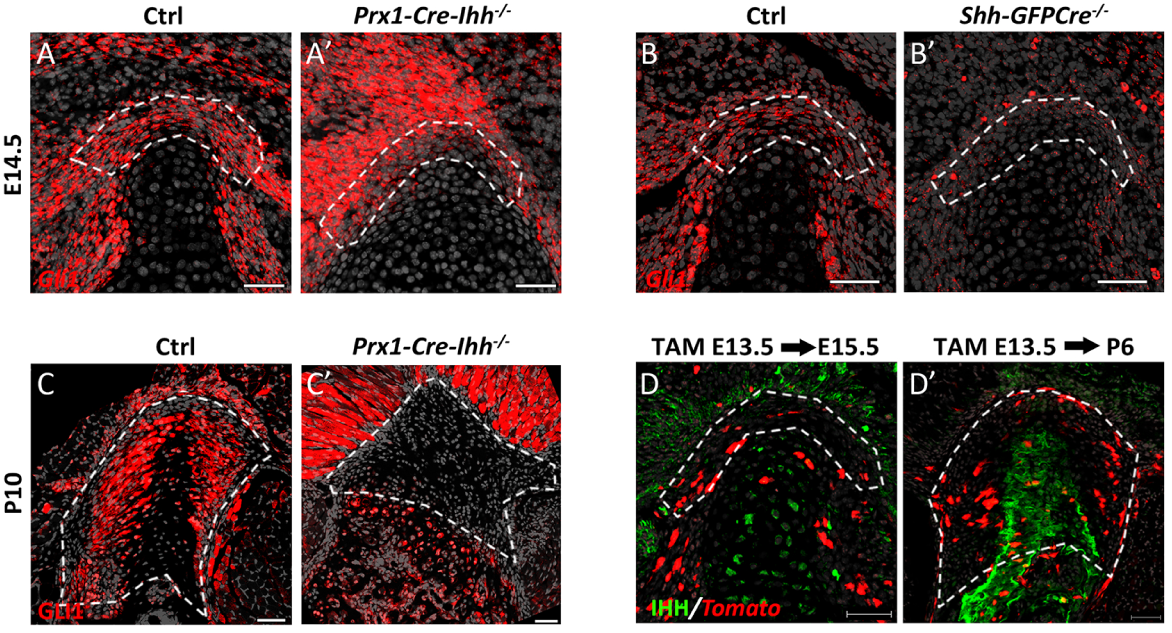
*Gli1* expression in migrating entheses is initiated by SHH and maintained by IHH. **(A,A’)** Fluorescent ISH for *Gli1* on sections through the DT of E14.5 control (A) and *Prx1-Cre-Ihh*^*f/f*^ embryos (A’) demonstrates that in the absence of IHH signaling, *Gli1* expression in the enthesis is unchanged. **(B,B’)** Fluorescent ISH for *Gli1* on sections through the DT of E14.5 control (B) and *Shh*^*Cre/Cre*^ embryos (B’) demonstrates that in the absence of SHH signaling, *Gli1* expression in the enthesis is lost. **(C,C’)** Immunostaining for GLI1 protein in *Prx1-Cre-Ihh*^*f/f*^ mutant (C’) and control (C) DT sections shows that in the absence of IHH, GLI1 protein expression is lost from the DT enthesis. **(D,D’)** Immunostaining for IHH protein on *Gli1-CreER^T2^;R26R-tdTomato* mice. Tamoxifen was administered at E13.5 and mice were sacrificed at E15.5 (D) or P6 (D’). IHH expression is seen in hypertrophic chondrocytes at E15.5 and in the mineralizing part of the enthesis at P6. Enthesis is demarcated by a dashed line. Scale bars: 50 µm.

Together, these results indicate that although *Gli1* lineage is not induced at the onset of enthesis formation (Blitz et al., 2013; Sugimoto et al., 2013), it contributes differentially to different entheses during embryonic development.

### *Gli1* expression in migrating entheses is initiated by SHH and maintained by IHH

The Hedgehog (HH) signaling pathway has previously been suggested to play a role in regulating the activity of *Gli1*-positive enthesis cells (Breidenbach et al., 2015; Dyment et al., 2015; Liu et al., 2013; Schwartz et al., 2015). Thus, our finding that *Gli1* expression is initiated at early stages of enthesis development (Fig. 1C(h)D(h)) raised the question of which component of the HH pathway regulates *Gli1* expression in enthesis cells and whether it also affects migratory entheses. It was suggested that Indian hedgehog (IHH), an effector of the HH signaling pathway, is a possible regulator of *Gli1* lineage cells in adult stationary entheses. In order to examine the possible role of IHH in inducing *Gli1* expression in the embryonic enthesis, we blocked *Ihh* expression in the limb by using *Prx1*-*Cre*-*Ihh*^*−/-*^ mice. Interestingly, at E14.5 *Gli1* was expressed in control and mutant entheses, suggesting that *Gli1* expression in embryonic enthesis progenitors is not regulated by *Ihh* (Fig. 3A,A’). We therefore examined the possible role of another regulator of HH signaling, namely sonic hedgehog (SHH), by analyzing *Gli1* expression by embryonic enthesis cells in *Shh* KO embryos (*Shh-GFPCre^−/-^*). Results showed that *Gli1* expression dramatically decreased in the mutant entheses compared to the WT (Fig. 3B,B’), suggesting that SHH is necessary for the induction of *Gli1* in embryonic enthesis cells.

That result was intriguing, because we observed that *Gli1* was constantly expressed by both embryonic and postnatal enthesis cells (Fig. 1A(h-h’’’),B(h-h’’’)), while *Shh* is expressed in the limb only during embryogenesis (Harfe et al., 2004). This raised the question of how *Gli1* expression in the enthesis is maintained after the loss of *Shh* expression. To address the possibility that *Ihh* controls *Gli1* expression in the postnatal enthesis even though it is not involved in the induction of *Gli1* expression in embryonic enthesis progenitors, we analyzed *Prx1*-*Cre*-*Ihh*^*−/-*^ mice at P10. Results showed that although *Gli1* expression was maintained in mutant muscle and bone, in the enthesis it was completely lost (Fig. 3C,C’), indicating that *Ihh* is indeed required for the maintenance of *Gli1* expression in enthesis cells. To identify the source of *Ihh* in the enthesis, we performed immunofluorescence staining for IHH protein in P6 enthesis sections. As seen in Figure 3 (D,D’), IHH was highly expressed in mineralized fibrocartilage and bone regions at the enthesis center.

Taken together, these results suggest that *Gli1* expression by embryonic enthesis progenitors is induced by SHH and later, in the maturing enthesis, maintained by IHH originating in mineralized fibrocartilage and bone.

### *Sox9* lineage cells of the embryonic enthesis do not contribute to postnatal migrating entheses

The embryonic enthesis originates from *Scx*/*Sox9* double-positive progenitor cells (Blitz et al., 2013; Sugimoto et al., 2013). Yet, the contribution of these progenitors to the postnatal enthesis and their relation to the *Gli1* lineage cells have never been studied. To fill this gap, we first examined the contribution of *Sox9* lineage to the postnatal enthesis. To that end, we performed a pulse-chase cell lineage experiment using mice that express Cre-ER under control of the *Sox9* promoter (*Sox9*-*CreER^T2^*) crossed with *R26R-tdTomato* reporter mice (Soeda et al., 2010). It was previously demonstrated that tamoxifen administration at E12.5 effectively labels embryonic enthesis cells (Blitz et al., 2013; Soeda et al., 2010). Indeed, examination at P0 following tamoxifen administration at E12.5 showed that the DT, TM and Achilles entheses were populated by tdTomato-positive cells, suggesting that at that stage, the enthesis is populated by *Sox9* lineage cells (Fig. 4A(a,b,d)). Yet, surprisingly, at P14 we observed a dramatic decrease in the contribution of tdTomato-expressing cells to the two migrating entheses (Fig. 4A(a’,b’,c), although the cells in the stationary enthesis were still extensively labeled (Fig. 4A(d’). These results suggest that postnatal stationary enthesis cells were descendants of the *Sox9*-positive embryonic lineage, whereas in migrating entheses this lineage was lost.

**Figure 4.**
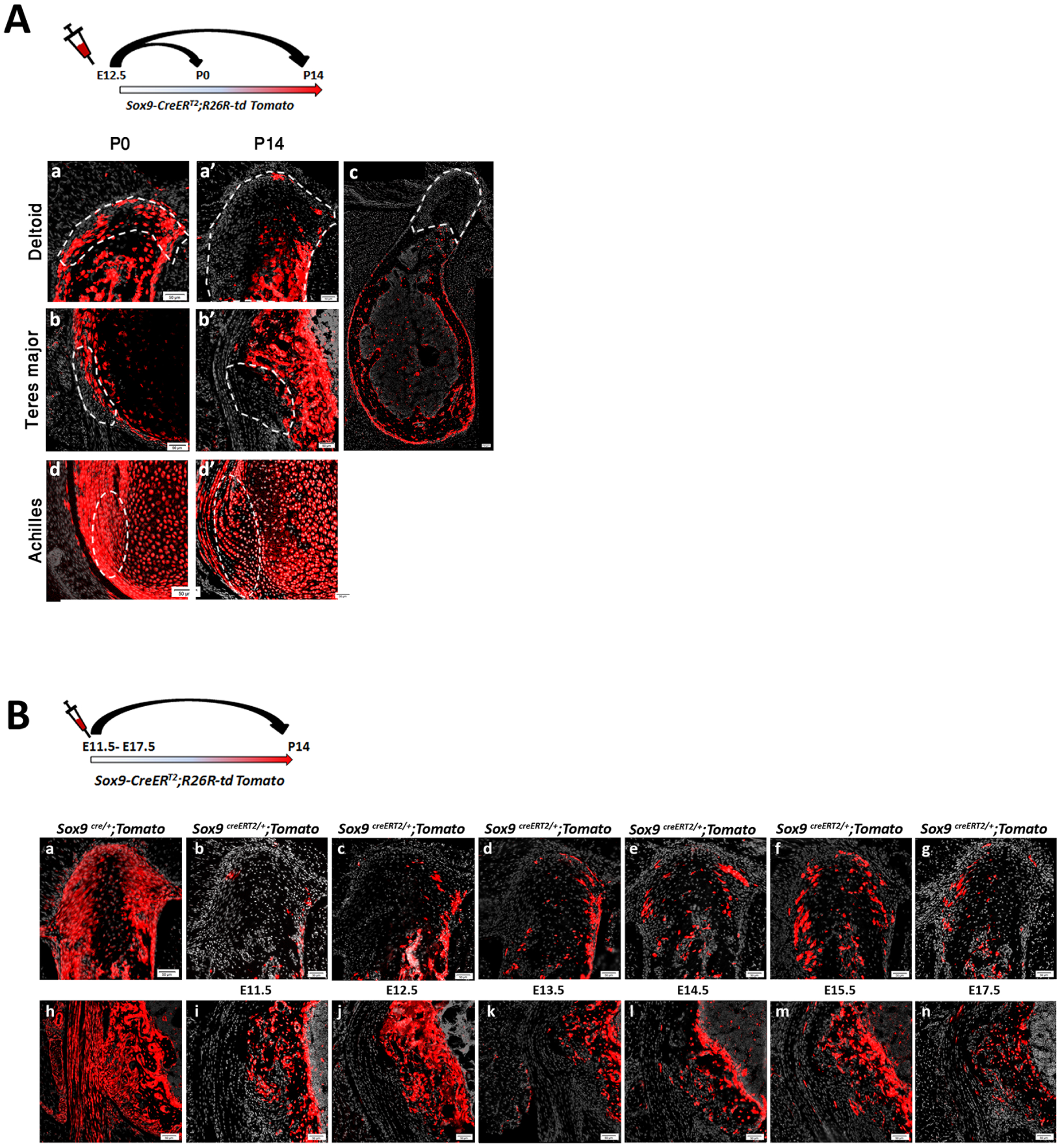
*Sox9* lineage cells of the embryonic enthesis do not contribute to postnatal migrating entheses. **(A)** Pulse-chase cell lineage experiment using *Sox9-CreER^T2^;R26R-td Tomato* mice demonstrates that *Sox9* lineage contributes differently to migrating and stationary entheses. *Sox9*-positive cells were marked at E12.5 by a single tamoxifen administration. At P0, tdTomato-positive cells were identified in migrating DT and TM entheses (A(a,b)) as well as in stationary Achilles entheses (A(d)). At P14, only a few tdTomato-positive cells were identified in migrating entheses (A(a’,b’,c)), whereas extensive staining was still seen in stationary entheses (A(d’)). Enthesis is demarcated by a dashed line. Scale bars: 50 µm. **(B)** The contribution of *Sox9* lineage to the DT and TM entheses was evaluated by crossing *Sox9-Cre* or *Sox9-CreER*^*T2*^ mice with *R26R-tdTomato* reporter mice. (Ba,h): The total contribution of the lineage was evaluated by examination of *Sox9-Cre;R26R-tdTomato* mice at P14. (Bb-g,i-n): The relative contribution of *Sox9* lineage cells specified at different time points was evaluated by pulse-chase experiments on *Sox9-CreER^T2^;R26R-tdTomato* mice, in which a single dose of tamoxifen was administered at various stages from E11.5 through E17.5.

The labeled stationary entheses could serve as a positive internal control for the effectiveness of labeling. Nevertheless, to rule out the possibility that tamoxifen was administered at the wrong time point, we repeated the experiment while administering tamoxifen at different time points from E11.5 to E17.5 (Fig. 4B). Examination at P14 showed minimal contribution of *Sox9* lineage cells to the postnatal entheses, similar to the results obtained following pulsing at E12.5. Taken together, these results indicate that the embryonic *Sox9* lineage contributes poorly to postnatal migrating entheses, suggesting that these entheses are populated by another cell lineage postnatally.

### *Gli1* lineage cells replace the embryonic *Sox9* lineage during enthesis maturation

Our finding that both embryonic *Gli1* and *Sox9* lineages contribute to the stationary postnatal enthesis suggests that these two genes mark a common cell lineage. Conversely, the finding that embryonic *Sox9* lineage contributes to embryonic but not to postnatal migrating enthesis implies that postnatally, embryonic *Gli1* lineage replaces the *Sox9* lineage to form the mature enthesis.

To study the process of lineage replacement and to follow its dynamics, we traced both lineages throughout enthesis development, from E15.5 to maturation at P14, by pulse-chase experiments. As seen in Figure 5, at E15.5 and P0 in both the DT and TM, *Sox9* lineage cells populated the embryonic enthesis. However, from P0 their number decreased dramatically and by P8, the enthesis contained only a limited number of these cells. Concurrently, the number of cells of the *Gli1* lineage gradually increased in both entheses and, by P8, *Gli1*+ cells inhabited most of the enthesis structure. Interestingly, the gradual population of the DT and TM entheses by *Gli1* lineage cells correlated with the temporal dynamics of enthesis migration (Fig. 1A). These results support our hypothesis that during early postnatal development of migrating entheses, a new population derived from *Gli1-*positive progenitors substitutes the *Sox9* lineage cells of the embryonic enthesis.

**Figure 5.**
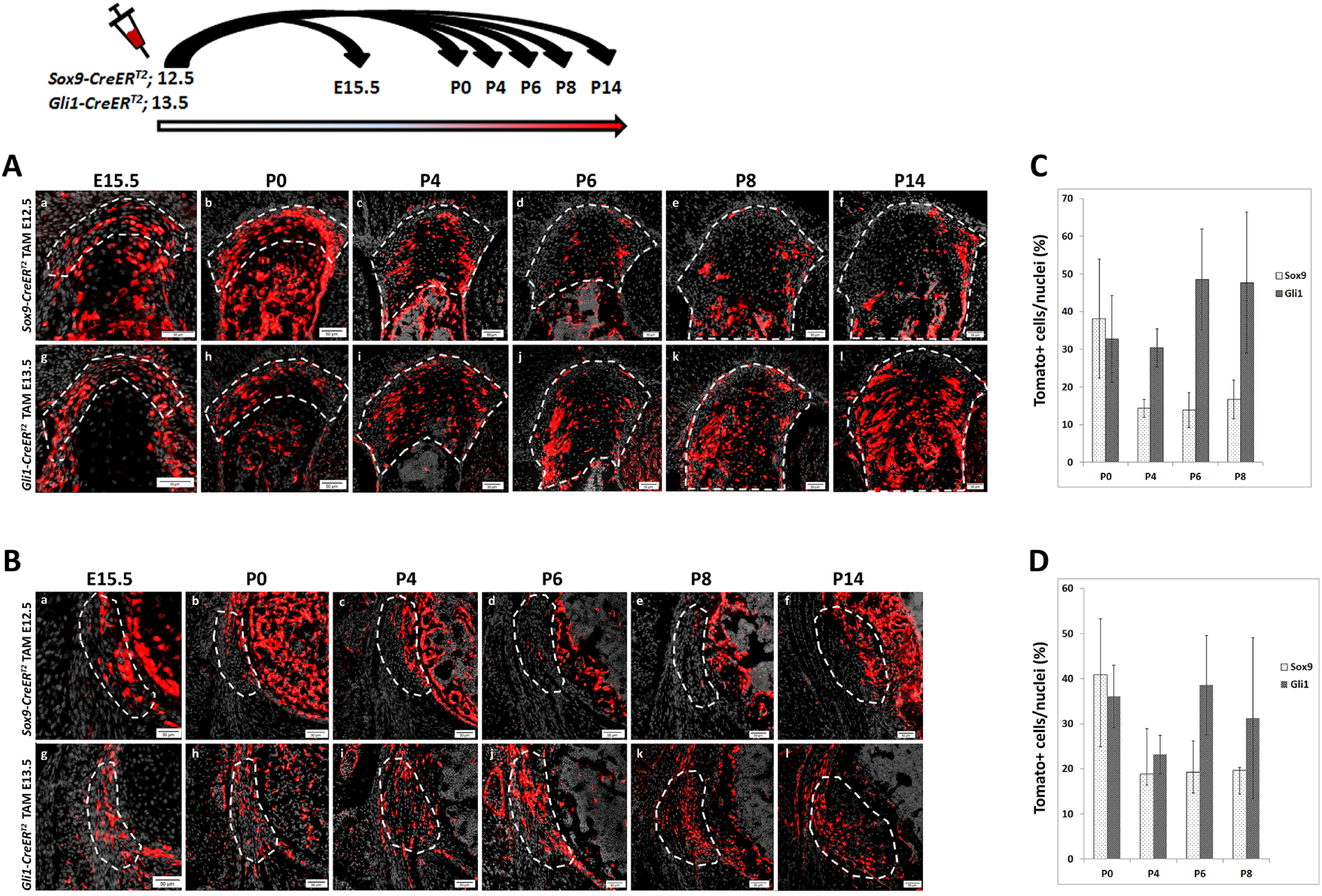
*Gli1* lineage cells replace *Sox9* lineage cells during enthesis maturation. **(A,B)** *Sox9-CreER^T2^;R26R-tdTomato* and *Gli1-CreER^T2^;R26R-tdTomato* mice were labeled by tamoxifen administration at E12.5 and E13.5, respectively and sacrificed at E15.5-P14. Transverse sections through the DT (A) and TM (B) show an increase in *Sox9* lineage cells at E15.5-P0. Between P0 and P14, a continuous decrease in *Sox9* lineage cells is seen. Concurrently, *Gli1* lineage cell number gradually increased in both DT and TM entheses. By P8, both entheses were populated by *Gli1* lineage cells. Enthesis is demarcated by a dashed line. Scale bars: 50 µm. **(C,D)** Graphs showing the percentage of *Gli1*-tdTomato and *Sox9*-tdTomato positive cells in P0, P4, P6, and P8 DT (C) and TM (D) cells.

### Cell fate of *Gli1*-positive progenitors is predetermined

The increasing cellular complexity of the developing enthesis and our finding that progenitors of the *Gli1* lineage that will contribute to the postnatal enthesis are present already during embryonic development led us to ask how these cells form the graded tissue of the mature enthesis. Specifically, we sought to determine whether *Gli1* lineage cells possess a multipotent capacity and, thereby, produce the different cell types of the various enthesis domains, or have predetermined cell fates already at the time of their recruitment to the embryonic enthesis. To decide between these options, we conducted pulse-chase experiments on *Gli1-CreER*^*T2*^ mice crossed with *R26R-Confetti* mice (Snippert et al., 2010) that, upon recombination, stochastically express GFP, RFP, CFP or YFP, allowing for the identification of different cell clones. Clones derived from multipotent progenitors would be spread among the different domains, whereas predetermined progenitors would produce clones that are confined to specific domains.

We focused on the two main domains of the DT enthesis, namely mineralized and non-mineralized fibrocartilage, and on the border between entheseal mesenchymal cells and bone in the TM enthesis. Examination of P14 limbs following tamoxifen administration at E13.5 revealed multiple clones in both entheses, most of which were restricted to a specific domain (Fig. 6). These results suggest that at E13.5, the cell fates of *Gli1* lineage progenitors have already been determined.

**Figure 6.**
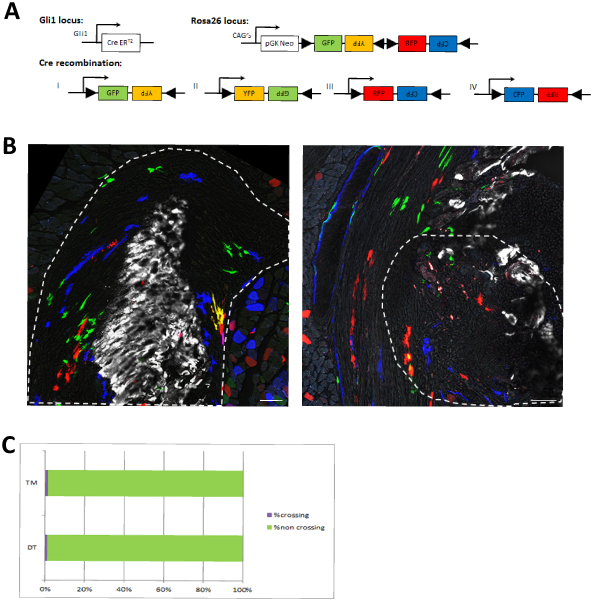
Cell fate of *Gli1*+ enthesis progenitors is pre-determined. Pulse-chase cell lineage experiments using *Gli1-CreER^T2^;R26R-Confetti* mice, in which *Gli1*+ progenitor cell clones were labeled. *Gli1*-positive cells were pulsed by a single tamoxifen administration at E13.5 and their descendants were followed to P14. One hour prior to sacrifice, mice were injected with Calcein blue. **(A)** Schematic illustration of possible Cre recombination outcomes. **(B)** Multiple clones were identified in the DT enthesis; however, the clones were restricted to either fibrocartilage or mineral fibrocartilage and did not cross the border between the layers. The clones identified in the TM enthesis were restricted to the fibrous part and did not penetrate the bone. (C) Graph showing the percentage of crossing vs. non-crossing clones. Scale bars: 50 µm.

## DISCUSSION

The transition from the embryonic to the mature enthesis has been understudied. Here, we use genetic lineage tracing to unravel the cellular developmental sequence of fibrous and fibrocartilaginous entheses. We show that although *Gli1*+ progenitors contribute to entheses of both types, their contribution differ greatly between stationary and migratory entheses. In stationary entheses, *Gli1*+ progenitors are descendants of *Sox9*+ progenitors that have established the embryonic enthesis. However, in migrating entheses, a separate, pre-specified population of *Gli1*+ progenitors is recruited to the embryonic enthesis, co-populates it alongside *Sox9*+ progenitors and, eventually, replaces them during postnatal development.

As mentioned, fibrous and fibrocartilaginous entheses exhibit marked differences in structure and composition (Benjamin et al., 2002). Notwithstanding the importance of this distinction, the observed differences in cellular origin between stationary and migrating entheses calls for a revision of the traditional classification and suggests that it should be taken into account whether an enthesis is “migratory” or “stationary”.

Organ development can be mediated by several cellular mechanisms. In a linear mechanism, an embryonic set of progenitors forms a primordium and then continues to proliferate and differentiate to form the mature organ. Another mechanism is cell recruitment, during which cells are supplied to the forming organ after primordium establishment. A third mechanism involves template replacement, in which an initial template is formed by one type of cells to be later replaced by another cell population, which will form the mature organ. Interestingly, in the musculoskeletal system there are examples of all these modes of development. Muscles and joints develop through cell recruitment (Buckingham et al., 2003; Shwartz et al., 2016), whereas most of the skeleton develops through template replacement, namely by endochondral ossification (Kronenberg, 2003). Here, we show that stationary entheses develop linearly by embryonic *Sox9*+ progenitors that form the postnatal enthesis and later upregulate the expression of *Gli1*. However, we show that migrating entheses develop through template replacement, as *Gli1*+ progenitors replace the *Sox9*+ embryonic enthesis cells and form the mature enthesis.

Previous studies have shown not only that *Gli1* is a marker for the forming enthesis, but that the HH pathway plays an active role in regulating enthesis maturation and regeneration (Breidenbach et al., 2015; Dyment et al., 2015; Liu et al., 2013; Schwartz et al., 2015; Schwartz et al., 2017). *Ihh* expression was identified in proximity to *Gli1*+ progenitor population in stationary entheses (Dyment et al., 2015; Liu et al., 2013; Schwartz et al., 2015). Moreover, loss-of-function of *Smo,* a componenet of the HH pathway, in *Scx*-expressing cells resulted in reduced mineralization of the enthesis (Breidenbach et al., 2015; Dyment et al., 2015; Liu et al., 2013; Schwartz et al., 2015). Yet, the question of which ligand of the HH pathway regulates the specification of *Gli1*+ cells in the embryo and, specifically, in migrating entheses has remained open. Our results clearly suggest that *Gli1* expression is regulated by both SHH and IHH during enthesis development. In the embryo, we show that SHH, but not IHH, signaling is essential for *Gli1* expression in the enthesis, implicating SHH in *Gli1*+ progenitor specification. However, during postnatal development, *Ihh* expression is vital for maintaining *Gli1* enthesis expression (Fig. 7A).

**Figure 7.**
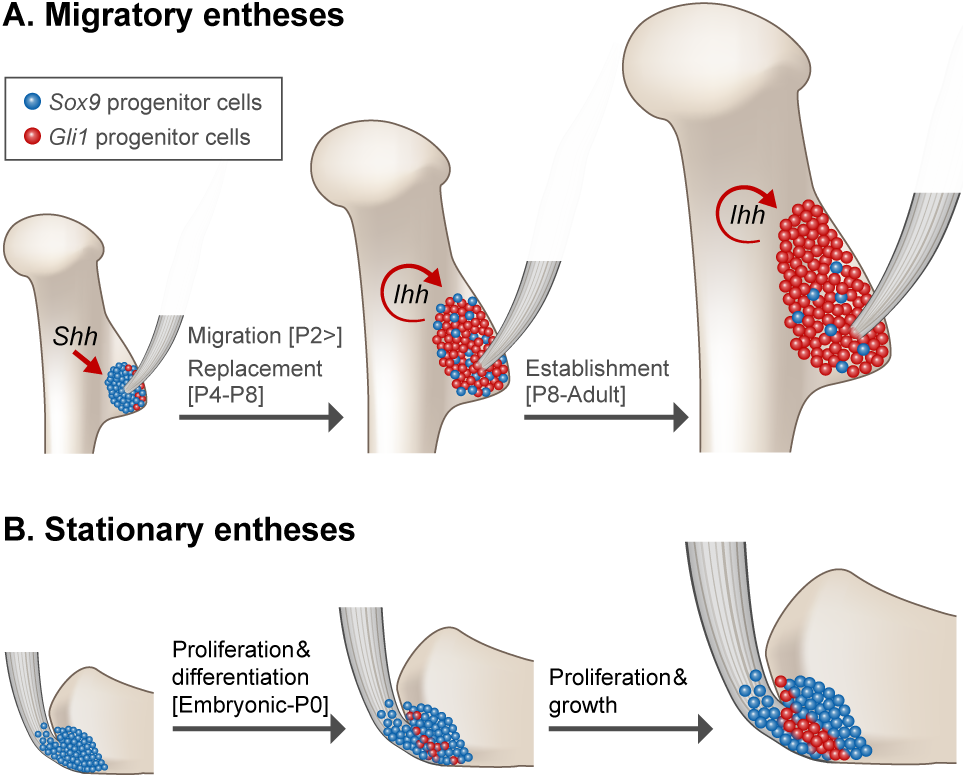
Migrating entheses develop through template replacement. Schematic illustration of the developmental sequence of migratory and stationary entheses. During embryogenesis, *Sox9* lineage cells form an enthesis template. **(A)** In migrating entheses, *Sox9* lineage cells are replaced by a second population of *Gli1* lineage cells, which eventually form the mature enthesis. Specification of *Gli1*lineage is regulated by SHH, whereas maintenance of this population is regulated by IHH. (B) In stationary entheses, embryonic *Sox9* lineage cells proliferate and differentiate and, eventually, populate the postnatal enthesis structure. During late embryonic development, some *Sox9* lineage cells start expressing *Gli1*.

During endochondral ossification, IHH regulates hypertrophic chondrocyte differentiation and, thereby, bone elongation (Vortkamp et al., 1996). It is therefore possible that by controling both these processes, IHH coordinates enthesis migration with the concurrent bone elongation to preserve enthesis position along the shaft. However, the signals that govern enthesis positioning and migration along the bone, including the possible effect of IHH on these processes, are yet to be elucidated.

Another key question regards the mechanism that facilitates the replacement between *Sox9*+ and *Gli1*+ cell populations. It is possible that the former cells die, while the latter cells proliferate and populate the entire enthesis. A more attractive hypothesis is that during enthesis drift, *Sox9*+ cells are removed together with eroded bone tissue by osteoclasts or other phagocytic cells. However, the replacement mechanism remains an open question. Moreover, whether the replacement process is the mechanism that allows migrating entheses to maintain their relative position along the bone is yet to be determined. Finally, our finding that *Gli1*-expressing cells are pre-determined already in the embryo suggests the existence of an earlier, yet unknown player in the specification of these progenitors. The discovery of this early marker gene may enable the identification of multipotent enthesis cells, which may hold therapeutic potential.

To conclude, our findings shed light on developmental linearity in organogenesis. Using the enthesis as a model system, we identify two different strategies of development (Fig. 7). The first, found in stationary entheses, is linear development where embryonic progenitors and their descendants contribute to the entire process of enthesis development, from embryogenesis to maturity. Conversely, in migrating entheses, another developmental strategy was identified where one type of progenitor cells form an embryonic template, only to be later replaced by another cell lineage that contributes to the mature organ.

## MATERIALS AND METHODS

### Animals

All experiments involving mice were approved by the Institutional Animal Care and Use Committee (IACUC) of the Weizmann Institute. Histology was performed on BL6 mice.

For lineage tracing experiments, *Sox9-Cre* (Akiyama et al., 2005), *Sox9-CreER* (Soeda et al., 2010) and *Gli1-CreER*^*T2*^ (Ahn and Joyner, 2004, Jackson laboratories) mice were crossed with *R26R-tdTomato* mice (B6;129S6-*Gt(ROSA)26Sor^tm9(CAG-tdTomato)Hze^*/J, (Madisen et al., 2010) or with *R26R-Confetti* mice (B6.129P2-*Gt(ROSA)26Sor^tm1(CAG-Brainbow2.1)Cle^*/J, Snippert et al., 2010).

To create *Shh* KO mice, mice heterozygous for a mutation in *Shh* (B6.Cg-*Shh^tm1(EGFP/cre)Cjt^*/J; Jackson Laboratory) were intercrossed; heterozygotes or WT littermates were used as a control. The generation of *Prx1*-*Cre* (Logan et al., 2002) was previously described. To generate *Prx1*-*Cre*-*Ihh* mutant mice, *Ihh*-floxed mice (B6N;129S4-*Ihh^tm1Blan^*/J; Jackson Laboratory) (Razzaque et al., 2005) were mated with *Prx1*-*Cre* mice. As a control, *Prx1*-*Cre*-negative animals were used.

For cell lineage experiments, *Sox9-CreER* ^*T2/+*^ or *Gli1-CreER*^*T2/+*^ males were crossed with *R26R-tdTomato* females to produce embryos carrying both the relevant *CreER*^*T2*^ and *R26R-tdTomato* alleles. For fate mapping, *Gli1-CreER*^*T2/+*^ males were crossed with *R26R-Confetti* females to produce embryos carrying both the *Gli1-CreER*^*T2*^ and *Confetti* alleles.

In all timed pregnancies, plug date was defined as E0.5. For harvesting of embryos, timed-pregnant females were sacrificed by cervical dislocation. The gravid uterus was dissected out and suspended in a bath of cold phosphate-buffered saline (PBS) and the embryos were harvested after removal of amnion and placenta. Tail genomic DNA was used for genotyping.

### Histological analysis, in situ hybridization and immunofluorescence

For histology and *in situ* hybridization, embryos were harvested at various ages, dissected, and fixed in 4% paraformaldehyde (PFA)/PBS at 4°C overnight. After fixation, tissues were dehydrated to 70% EtOH and embedded in paraffin. The embedded tissues were cut to generate 7-µm-thick sections and mounted onto slides. Hematoxylin and eosin (H&E) staining was performed following standard protocols. Non-fluorescent and fluorescent *in situ* hybridizations were performed as previously described using digoxigenin-(DIG) labeled probes (Shwartz and Zelzer, 2014). All probes are available upon request.

For immunofluorescence staining, 10-µm-thick cryosections were air-dried for 1 hour before staining. For IHH staining, sections were washed twice in PBST for 5 minutes and blocked to prevent non-specific binding with 7% goat serum and 1% BSA dissolved in PBST. Then, sections were incubated with rabbit anti-IHH antibody (Abcam, #AB39364, 1:50) at 4C° overnight. The next day, sections were washed twice with PBST and incubated for 1 hour with Cy2 conjugated fluorescent antibody (1:100, Jackson Laboratories). Slides were mounted with Immuno-mount aqueous-based mounting medium (Thermo).

For Gli1 staining, 10-µm-thick cryosections were air dried for 1 hour and fixed in 4% PFA for 10 minutes. Then, sections were washed twice in PBST and endogenous peroxidase was quenched using 3% H_2_O_2_ in PBS. Next, antigen retrieval was preformed using 0.3% Triton in PBS. Non-specific binding was blocked using 7% horse serum and 1% BSA dissolved in PBST for 1 hour. Then, sections were incubated with Goat anti-GLI1 antibody (R&D systems, #AF3455, 1:100) overnight at room temperature. The next day, sections were washed twice in PBST and incubated with Biotin anti-goat (1:100, Jackson laboratories, 705065147) for 1 hour and then with streptavidin-HRP (1:200 Perkin Elmer, NEL750001EA) for 1 hour. HRP was developed using TSA amplification kit (1:100, Prekin Elmer) for 20 minutes, counterstained with DAPI and mounted with Immuno-mount aqueous-based mounting medium (Thermo).

### Physical position of entheses

For identification of enthesis physical position, 2-4 limb samples at stages E16.5-P14 were scanned *ex vivo* using iodine contrast agent to allow visualization of the soft tissue. DT and TM positions were identified manually, and their physical position along the bone was calculated as described previously (Stern et al., 2015).

### Cell lineage analysis

Tamoxifen (Sigma T-5648) was dissolved in corn oil (Sigma C-8267) at a final concentration of 5 mg/ml. Time-mated *R26R-tdTomato* females were administered 1 mg of tamoxifen by oral gavage (FST) at designated time points as indicated.

### Cell count

For each age, 2-4 limbs from different litters were harvested, embedded in OCT, and sectioned at a thickness of 10 µm. Sections were imaged using Zeiss LSM 780 microscope approximately every 80 µm, capturing red and DAPI channels. Images were then processed in ImageJ as follows: Each image was converted to an RGB stack and the region of interest was manually identified and cropped. Then, image levels were adjusted to improve separation between nuclei. A binary threshold was set automatically and conjoined nuclei were automatically separated using the binary watershed function. Using a home-made Matlab script, the number of nuclei was counted. Each nucleus that was co-localized with a red channel signal was counted as a tdTomato-positive cell. For each section, the percentage of tdTomato-positive cells was calculated. The average percentage of these cells was calculated first for each bone and then for all samples. From each line and for each age group, at least 3 individual bones were sampled, and 9-17 sections per bone were analyzed, depending on bone length. Data are presented as mean ± SD.

### Fate mapping Confetti experiment

Tamoxifen (Sigma T-5648) was dissolved in corn oil (Sigma C-8267) at a final concentration of 20 mg/ml. Time-mated *R26R-Confetti* females were administered 4 mg of tamoxifen by oral gavage at E13.5. Cre-positive pups were sacrificed at P14. 1 hour prior to sacrifice, each pup was injected with Calcein blue (Sigma, #m1255; 30 mg/kg). Limbs were harvested, fixed for 30 minutes in 4% PFA at 4°C, embedded in OCT, and sectioned at a thickness of 10 µm. Sections were imaged using Zeiss LSM 780 microscope approximately every 80 µm.

### Microscope settings and image analysis

At least 1024×1024 pixels, 8-bit images were acquired using the X20 lens. A Z-stack of 2-4 images was taken from each section. To detect GFP and YFP, the argon laser 488 nm was used. For RFP detection, a red diode laser emitting at 561 nm was used, and blue mCFP was excited using a laser line at 458 nm. Calcein blue staining was detected using the 405-nm laser line. GFP fluorescence was collected between ~500-598 nm, airy 1; RFP fluorescence was collected between ~606-654nm, airy 1; mCFP fluorescence was collected between ~464-500 nm, airy 1 and Calcein blue was collected between ~410-451. For each image, a corresponding bright field image was captured. The acquired images were processed using Photoshop and ImageJ.

To calculate the percentage of *Gli1*+ clones found in both mineralized and non-mineralized fibrocartilage, the border between the zones was identified by Calcein blue signal. Clones that crossed the mineral border were manually counted and their percentage was calculated first for each section, then for each limb and, finally, for each type of enthesis. Data are presented as mean ± SD.

## Acknowledgments

We thank Nitzan Konstantin for expert editorial assistance. We thank Dr. Patrick Tschopp from the Clifford J. Tabin lab, Harvard Medical School, for his assistance in generating *Shh* KO mice. Special thanks to all members of the Zelzer laboratory for encouragement and advice.

This study was supported by grants from the National Institutes of Health (grant #R01 AR055580), European Research Council (ERC) (grant #310098), the Jeanne and Joseph Nissim Foundation for Life Sciences Research, the Y. Leon Benoziyo Institute for Molecular Medicine, Beth Rom-Rymer, the Estate of David Levinson, the Jaffe Bernard and Audrey Foundation, Georges Lustgarten Cancer Research Fund, the David and Fela Shapell Family Center for Genetic Disorders, the David and Fela Shapell Family Foundation INCPM Fund for Preclinical Studies, and the Estate of Bernard Bishin for the WIS-Clalit Program.

